# Disruption of Fuz in mouse embryos generates hypoplastic hindbrain development and reduced cranial nerve ganglia

**DOI:** 10.1101/2023.08.04.552068

**Authors:** Carlo Donato Caiaffa, Yogeshwari S. Ambekar, Manmohan Singh, Ying Linda Lin, Bogdan Wlodarczyk, Salavat R. Aglyamov, Giuliano Scarcelli, Kirill V. Larin, Richard Finnell

## Abstract

The formation of the brain and spinal cord is initiated in the earliest stages of mammalian pregnancy in a highly organized process known as neurulation. Convergent and extension movements transforms a flat sheet of ectodermal cells into a narrow and elongated line of neuroepithelia, while a major source of Sonic Hedgehog signaling from the notochord induces the overlying neuroepithelial cells to form two apposed neural folds. Afterward, neural tube closure occurs by synchronized coordination of the surface ectoderm and adjacent neuroepithelial walls at specific axial regions known as neuropores. Environmental or genetic interferences can impair neurulation resulting in neural tube defects. The *Fuz* gene encodes a subunit of the CPLANE complex, which is a macromolecular planar polarity effector required for ciliogenesis. Ablation of *Fuz* in mouse embryos results in exencephaly and spina bifida, including dysmorphic craniofacial structures due to defective cilia formation and impaired Sonic Hedgehog signaling. In this work, we demonstrate that knocking *Fuz* out during embryonic mouse development results in a hypoplastic hindbrain phenotype, displaying abnormal rhombomeres with reduced length and width. This phenotype is associated with persistent loss of ventral neuroepithelial stiffness, in a notochord adjacent area at the level of the rhombomere 5, preceding the development of exencephaly in *Fuz* ablated mutants. The formation of cranial and paravertebral ganglia is also impaired in these embryos, indicating that *Fuz* has a critical function sustaining normal neural tube development and neuronal differentiation.

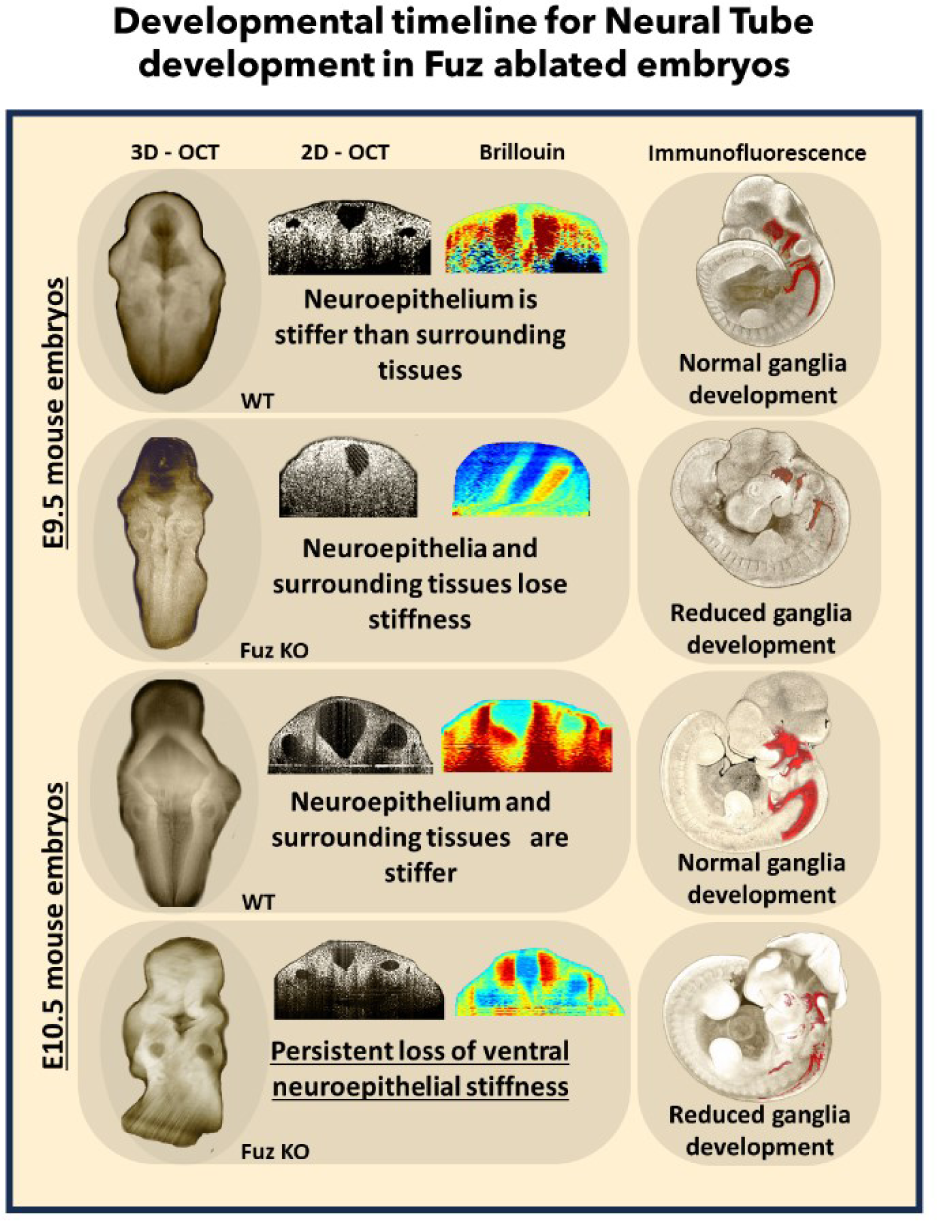

**SIGNIFICANCE STATEMENT:** Neural tube defects (NTDs) are a common cause of disability in children, representing the second most common congenital structural malformation in humans following only congenital cardiovascular malformations. NTDs affect approximately 1 to 2 pregnancies per 1000 births every year worldwide, when the mechanical forces folding the neural plate fails to close at specific neuropores located anteriorly (cranial) or posteriorly (caudal) along the neural tube, in a process known as neurulation, which happens throughout the third and fourth weeks of human pregnancy.

## INTRODUCTION

The World Health Organization estimates that every year 8 million babies are born worldwide with a birth defect. Cardiac defects, neural tube defects (NTDs) and Down syndrome are reported as the most common severe birth defects, which are primarily caused by an either genetic mutations, maternal infections, nutritional imbalances, environmental factors, or an interaction between two or more of these factors (Perin *et al*., 2023; Finnell *et al*.2021; Caiaffa *et al*.,2023).

NTDs are a consequence of disturbances during neurulation, which is one of the earliest and most important ontogenetic milestones, set right before organogenesis, and continuing until the embryonic structures of the brain and spinal cord are developed (Copp *et al*., 2003). The genetic network required for neurulation has been rigorously investigated in vertebrate models of embryonic development. From these studies, a total of 340 genes were identified for being associated with abnormal neural tube closure phenotype in different mouse models (MGI mouse phenotype annotation - MP:0003720). The vast majority of studied NTD mutants present with a neural tube phenotype including exencephaly (225 annotated genes, MP:0000914) and spina bifida (60 annotated genes, MP:0003054), while a minority of the described mutants can be characterized as having craniorachischisis (17 annotated genes, MP:0008784). Remarkably, the craniorachischisis phenotype observed in mice embryos is associated with genes coding for the “core” Planar Cell Polarity regulators (including *Celsr1; Dact1; Dvl1; Dvl2; Dvl3; Med12; Ptk7; Scrib; Sfrp1; Sfrp2; Sfrp5; Vangl1; Vangl2*).

The *Fuz* gene (*fuzzy planar cell polarity* gene) encodes a planar cell polarity effector protein required for ciliogenesis and the organization of directional cellular movement. Ablation of *Fuz* in mouse embryos generates several embryonic defects including defective cilia, abnormal neural tube closure and dysmorphic craniofacial structures, and variants in this gene are associated with NTDs and craniosynostosis in humans (Park *et al*., 2006; Gray *et al*., 2009; Seo *et al*., 2011; Barrell *et al*., 2022). Ablation of the *Fuz* gene in mouse embryos does not generate craniorachischisis, rather the embryos analyzed during organogenesis present with exencephaly and/or spina bifida, which are also found as known phenotypes for mutants in the Sonic Hedgehog (Shh) pathway and in mouse models for ciliopathies. In fact, considering both Shh mutants and ciliopathy models, there are 32 genes annotated on MGI for development of exencephaly including: *Fuz, Gli2, Gli3, Gpr161, Intu, Ptch1, Rab23, Sufu* and *Zic2*.

In vertebrates, the evolutionarily conserved abilities of the primary cilia for the transduction of signals that will modulate cellular behavior, perception, and motility, after receiving biomechanical and chemical extracellular stimuli, are fundamental requirements for normal development of several anatomical structures including the brain and the spinal cord (Echelard *et al*., 1993; Gray *et al*., 2009; Heydeck *et al*., 2009), inner ear (Jones and Chen, 2008), kidneys (Jonassen *et al*., 2008), larynx (Tabler *et al*., 2017) and the eyes (Fiori *et al*., 2020).

The FUZ protein physically interacts with INTU and WDPCP (Langousis *et al*., 2022), as individual monomeric subunits, to form the CPLANE macromolecular complex (ciliogenesis and planar polarity effector complex), which is required for cilia formation via regulation of apical actin assembly (Park *et al*., 2006), septin modulation (Kim *et al*., 2010), assembly of intraflagellar transport complex IFT-A (Toriyama *et al*., 2016) and binding to Rab23 and Rsg1 during the final steps in ciliogenesis (Agbu *et al*., 2018; Strutt *et al*., 2019). Genetic ablation experiments in the mouse revealed that both *Fuz* and *Intu* genes are required for neural tube and craniofacial formation. These genes are also critical in generating phenotypes linked to cilia defects, including polydactyly, skeletal defects, abnormal Sonic Hedgehog signaling and Gli3 processing (Zeng *et al*., 2010; Chang *et al*., 2015). Besides WDPCP participation as a core interaction partner of the constituted CPLANE complex, neural tube closure was not yet systematically studied in Wdpcp ablated embryos, although they do share a similar craniofacial phenotype, including facial and palate clefts accompanied by polydactyly, abnormal truncus arteriosus and atrioventricular septal defects, when compared with those described as *Fuz* and *Intu* mutant phenotypes (Cui *et al*., 2013).

Observations of the abnormal neural tube closure phenotypes reported in mouse mutant embryos for the CPLANE proteins and also for the downstream components of the Sonic Hedgehog pathway (*Gli3, Gpr161, Rab23* and *Sufu*), which are not linked to a holoprosencephaly phenotype (*Shh, Smo, Stil, Hhat, Cdon, Gas1* – upstream on Shh pathway), point to a direction where *Fuz, Intu*, and most probably, *Wdpcp*, are required during ciliogenesis to sustain proper transduction of the Sonic Hedgehog signaling pathway on neuroepithelial cells.

Fuz is a highly conserved gene, comprehensively studied in model organisms such as mice, Zebrafish and *Xenopus* (Grey et al., 2009; Heydek et al., 2009; Zhang et al., 2011; Seo et al., 2011). As the ablation of Fuz is not linked to craniorachischisis, the typical PCP mutant phenotype, rather been associated with exencephaly and an abnormal regulation of ciliogenesis and ciliary function, we planned to investigate how abnormal cilia development influence hindbrain morphogenesis and most importantly the neuroepithelial organization during neural tube closure.

## RESULTS

### Disruption of Fuz generates hypoplastic hindbrain and abnormal rhombomeres with diminished length and width

The most anterior region of the neural plate will establish the forebrain, midbrain, and hindbrain, respectively. The hindbrain is segmented into rhombomeres, that will guide the organization of neuronal projections throughout the adjacent craniofacial tissues (Krumlauf and Wilkinson, 2021). In this study, we observed the hindbrain development of *Fuz* ablated mouse embryos at E9.5 and E10.5, using both confocal microscopy and optical coherence tomography (OCT) images (Figure 1 a-i). Image quantification of the distances in between the rhombomere borders indicate that the hindbrain of *Fuz* mutant embryos do not develop normally, with abnormal rhombomeres due to their diminished length and width. Considering the most anterior even numbered rhombomeres, r2 and r4, this difference in the measured length, from left to right borders, can be detected as early as E9.5, and becomes more accentuated at E10.5 (Figure 1 - Rhombomere Length).

**Figure 1.**
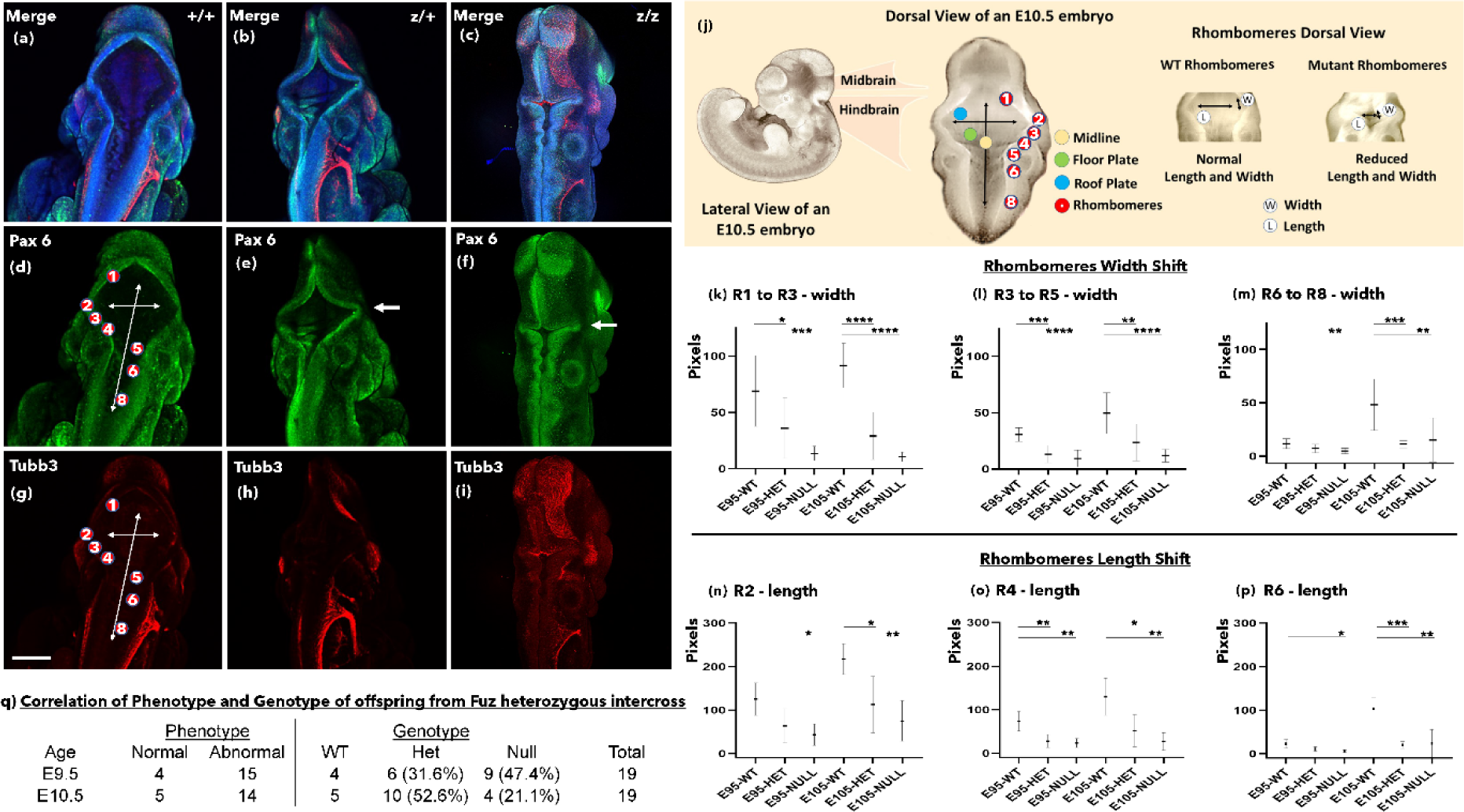
Fuz mutant embryos develop abnormal rhombomeres with diminished length and width. Confocal imaging representing the hindbrains of Fuz embryos at E10.5. (a, d, g) merged channels; (b, e, h) Pax6 immunostaining; (c, f, i) Tubb3 immunostaining. (j) Illustration of the mouse embryo hindbrain showing the identification of wildtype and mutant rhombomeres. (k) Rhombomeres 1 to rhombomeres 3 width shift measured in pixels, (l) Rhombomeres 3 to rhombomeres 5 width shift (pixels), (m) Rhombomeres 6 to rhombomeres 8 width shift (pixels), (n) Rhombomeres 2 Length shift measured in pixels, (o) Rhombomeres 4 Length shift (pixels), Rhombomeres 6 Length shift (pixels), * and **** indicate P < 0.05 and P <0.0001, respectively by One-Way ANOVA and Bonferroni’s comparisons test. (q) Correlation of Phenotype and Genotype of offspring from Fuz heterozygous intercross. The scale bar is 0.5 mm.

The distance between boundaries of the odd numbered rhombomeres r1 to r3, and r3 to r5, and between the more posterior even numbered rhombomeres, r6 to r8, was also measured. The decreased boundary distance in between the odd rhombomeres r1 to r3 can be noticed at E9.5, while decreased boundary distances from r1 to r3, r3 to r5 and r6 to r8 becomes more accentuated at E10.5, indicating that the overall tissue structure of the hindbrain does not develop to its full potential in the absence of the Fuz gene product (Figure 1 - Rhombomere Width).

### Fuz mutant embryos exhibit persistent loss of ventral neural fold stiffness

In vertebrate embryos, neurons descend from three different lineages arising from the neural tube neuroepithelium, neural crest, and cranial placodes, all of which are derived from the ectodermal germinal layer (Gans and Northcutt, 1983; Briscoe *et al*., 2000; Koontz *et al*., 2023). Defective cilium impairs Shh signaling, and as a consequence, disruption of normal development of ventral neuroepithelial cells is observed during neurulation (Ma *et al*., 2002; Caspary *et al*., 2002; Huangfu *et al*., 2003; Martinez-Chavez *et al*., 2018).

We focused on measuring hindbrain neuroepithelial stiffness at the level of the rhombomere 5, in an area adjacent to the otic pits, a primordial embryonic vesicle derived from the ectodermal placodes, which give rise to the inner ear and is located in a hindbrain region that does not provide a significant contribution of neural crest cells to the mesenchyme (Trainor *et al*., 2002). Our stiffness measurements during neurulation indicate that tissue stiffness decreases in the neural folds at E9.5 in all of the mutants analyzed (Figure 2), while at E10.5 the dorsal areas of the neural folds partially recover stiffness, indicating a persistent loss of ventral neural fold stiffness throughout neural tube development in the Fuz ablated embryos (Figure 3).

**Figure 2.**
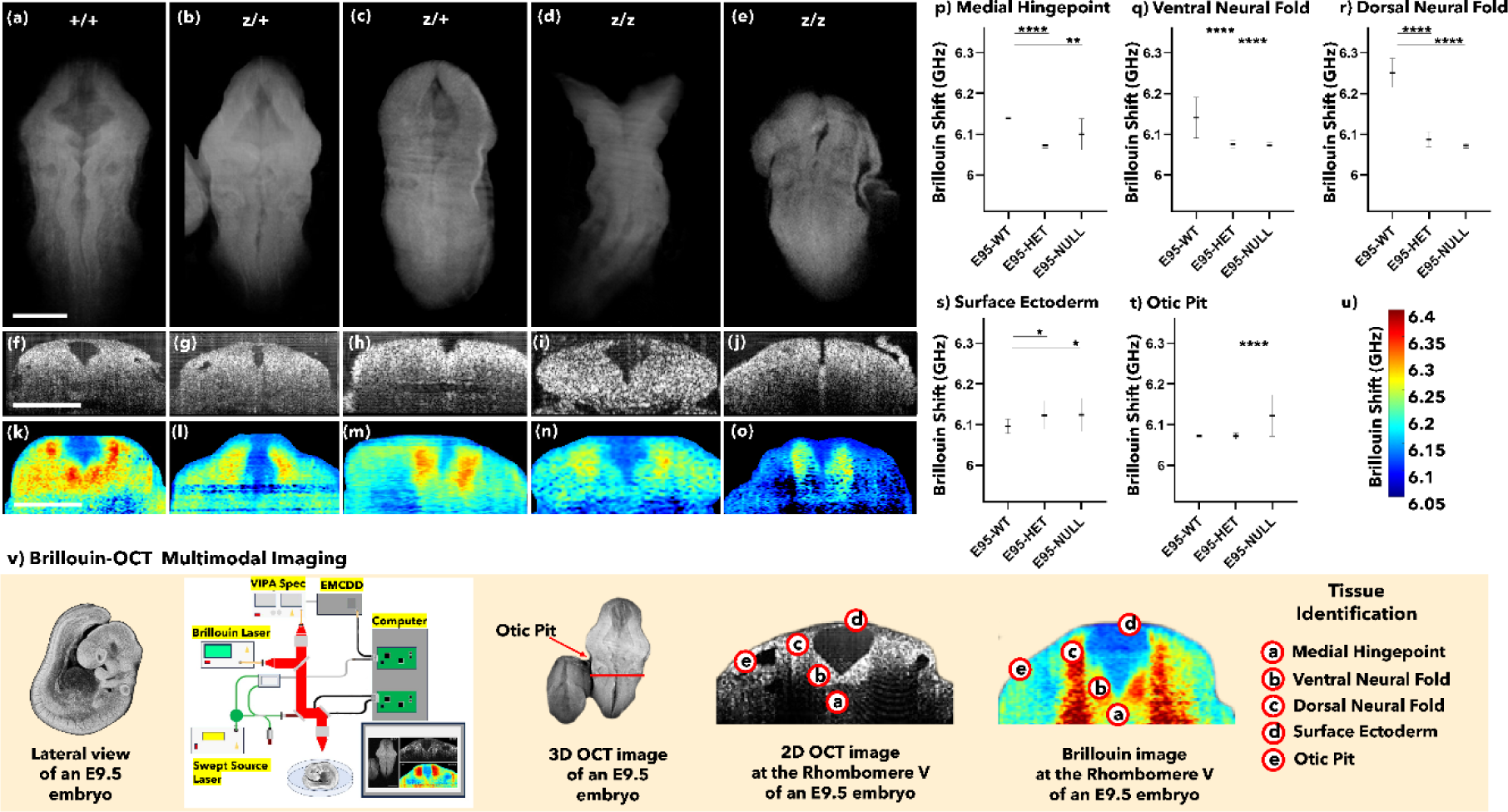
Fuz mutant embryos present with decreased neural fold stiffness at E9.5. 3D-OCT images are showing the hindbrains of Fuz embryos at E 9.5 (a-e). 2D-OCT optical sections are showing the neural folds at the rhombomere 5 (f-j). Brillouin frequency shift images represent tissue stiffness in the anatomical areas identified by 2D-OCT optical sections (k-o). Region-wise average Brillouin frequency shift of wild-type (WT), heterozygous (HET), and nullizygous (NULL) embryos at the (p) Medial Hingepoint, (q) Ventral Neural Fold, (r) Dorsal Neural Fold, (s) Surface Ectoderm and (t) Otic Pit regions, * and **** indicate P < 0.05 and P <0.0001, respectively by One-Way ANOVA and Bonferroni’s comparisons test. (u) Brillouin frequency shift Color Bar and (v) Illustration of the Experimental Scheme for Tissue Stiffness Measurements using the Brillouin-OCT Multimodal Imaging System. The scale bar is 0.25 mm.

**Figure 3.**
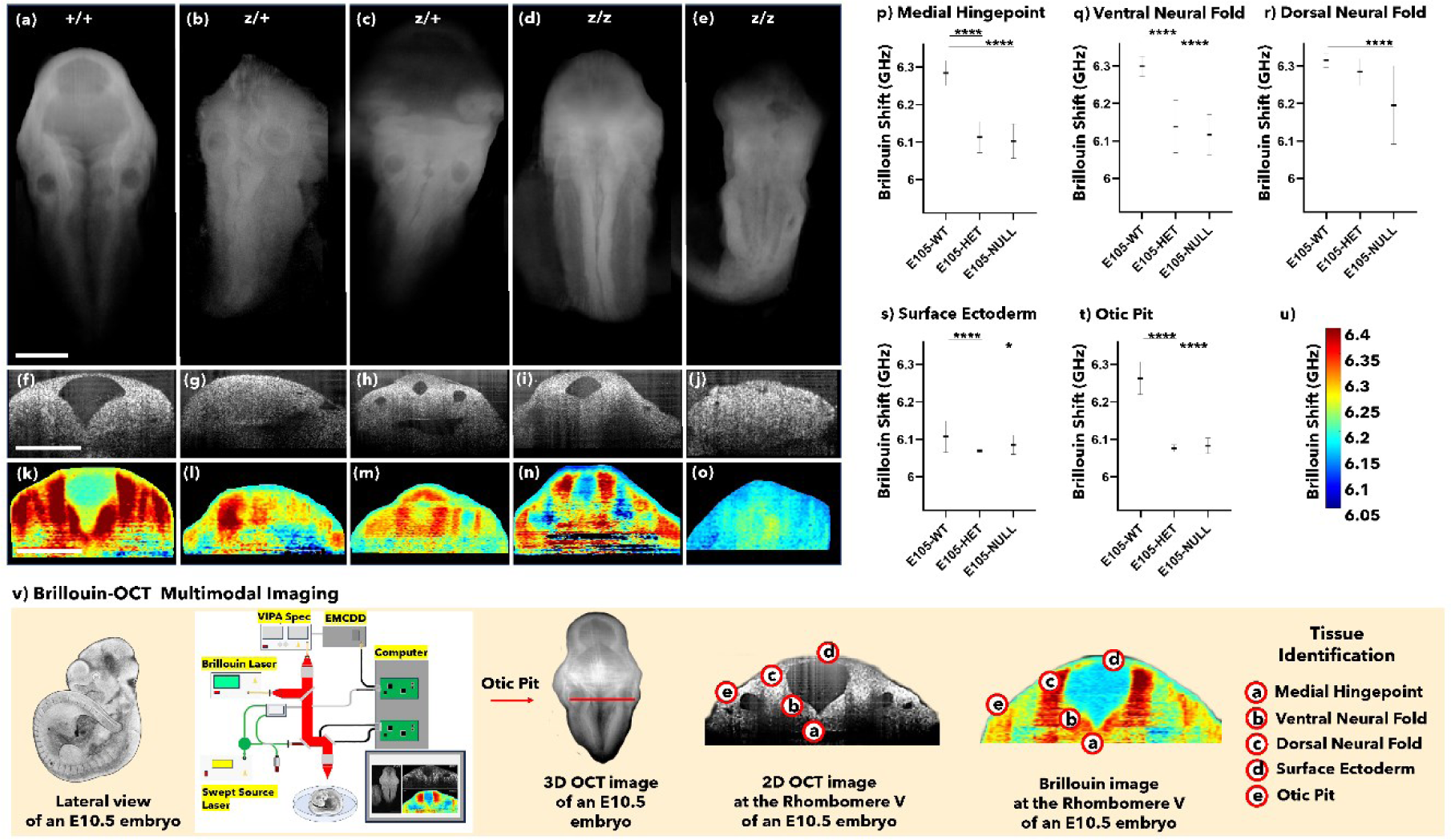
Fuz mutant embryos exhibit persistent loss of ventral neural fold stiffness at E10.5. 3D-OCT images are showing the hindbrains of Fuz embryos at E 10.5 (a-e). 2D-OCT optical sections are showing the neural folds at the rhombomere 5 (f-j). Brillouin frequency shift images represent tissue stiffness in the anatomical areas identified by 2D-OCT optical sections (k-o). Region-wise average Brillouin frequency shift of wild-type (WT), heterozygous (HET), and nullizygous (NULL) embryos at the (p) Medial Hingepoint, (q) Ventral Neural Fold, (r) Dorsal Neural Fold, (s) Surface Ectoderm and (t) Otic Pit regions, * and **** indicate P < 0.05 and P <0.0001, respectively by One-Way ANOVA and Bonferroni’s comparisons test. (u) Brillouin frequency shift Color Bar and (v) Illustration of the Experimental Scheme for Tissue Stiffness Measurements using the Brillouin-OCT Multimodal Imaging System. The scale bar is 0.5 mm.

### Cranial and paravertebral ganglia development are reduced in Fuz ablated embryos

Given the importance of Fuz during neuroepithelial development, we hypothesized that neuronal differentiation would decrease in Fuz ablated embryos. Therefore, the neural fate from Pax 6 positive derived cells would expect to be disrupted in the cranial ganglia and in posterior areas along the rostral caudal axis. In support of this idea, we performed immunostaining to detect Pax6 and Tubb3 (neurofilament - Tuj1 monoclonal serum). *Fuz* ablated embryos present with reduced cranial and paravertebral ganglia development, and the pattern of differentiated neurons along the body axis is significantly decreased (Figure 4), indicating that the function of Fuz in ciliogenesis is an important requirement preceding neurogenesis that originated from neural plate derived tissues.

**Figure 4.**
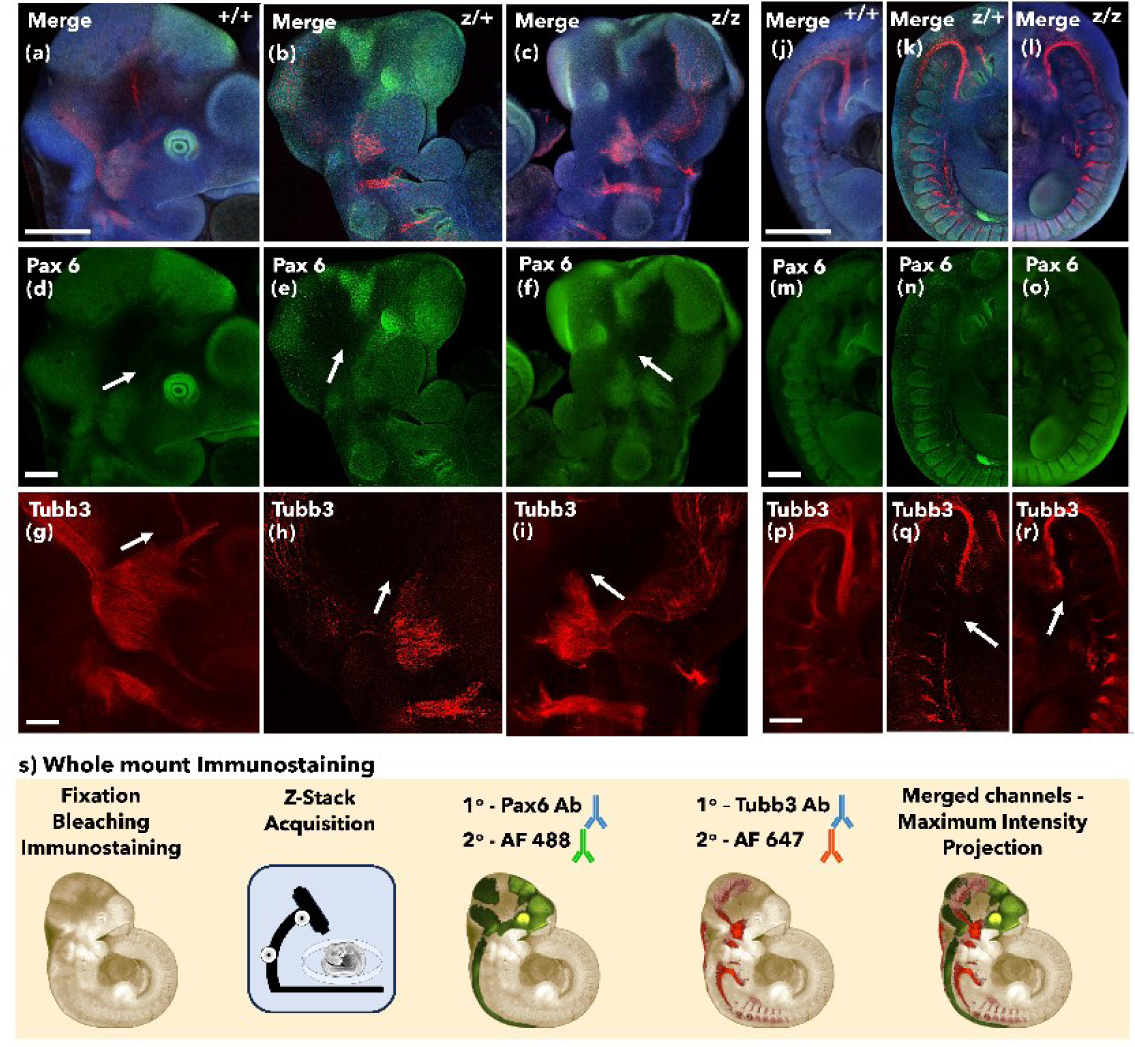
Fuz mutant embryos exhibit reduced cranial and paravertebral ganglia. Confocal imaging representing the cranial and caudal views of Fuz embryos at E10.5. (a, b, c) Cranial merged channels; (d, e, f) Cranial Pax6 immunostaining; (g, h, i) Cranial Tubb3 immunostaining; (j, k, l) Caudal merged channels; (m, n, o) Caudal Pax6 immunostaining; (p, q, r) Caudal Tubb3 immunostaining. (s) Illustration of the whole mount immunostaining protocol and image acquisition on a confocal microscope. The scale bar is 0.5 mm.

### *Fuz* ablated embryos exhibit abnormal Sox10 positive neural crest migration

The cranial ganglia including the trigeminal ganglia is derived from both cranial neural crest and ectodermal placodes (Steventon *et al*., 2014). After detecting the importance of Fuz during the development of the neural tube and cranial ganglia, we decided to investigate the migratory behavior of hindbrain derived neural crest cells in *Fuz* ablated embryos. We anticipated that the neural fate from Sox10 positive neural crest cells would be impaired in the cranial ganglia and in the posterior areas along the rostral caudal axis. In support of this idea, we performed immunostainings to detect Pax3 and Sox10 (Figure 5). The normal Pax3 staining pattern observed in the wildtype is disrupted at the midbrain-hindbrain boundary in the Fuz ablated embryos. We also observed an abnormal Sox10 positive neural crest cell migration pattern in an area adjacent to the first pharyngeal arch, where these cells accumulate during trigeminal ganglia development.

**Figure 5.**
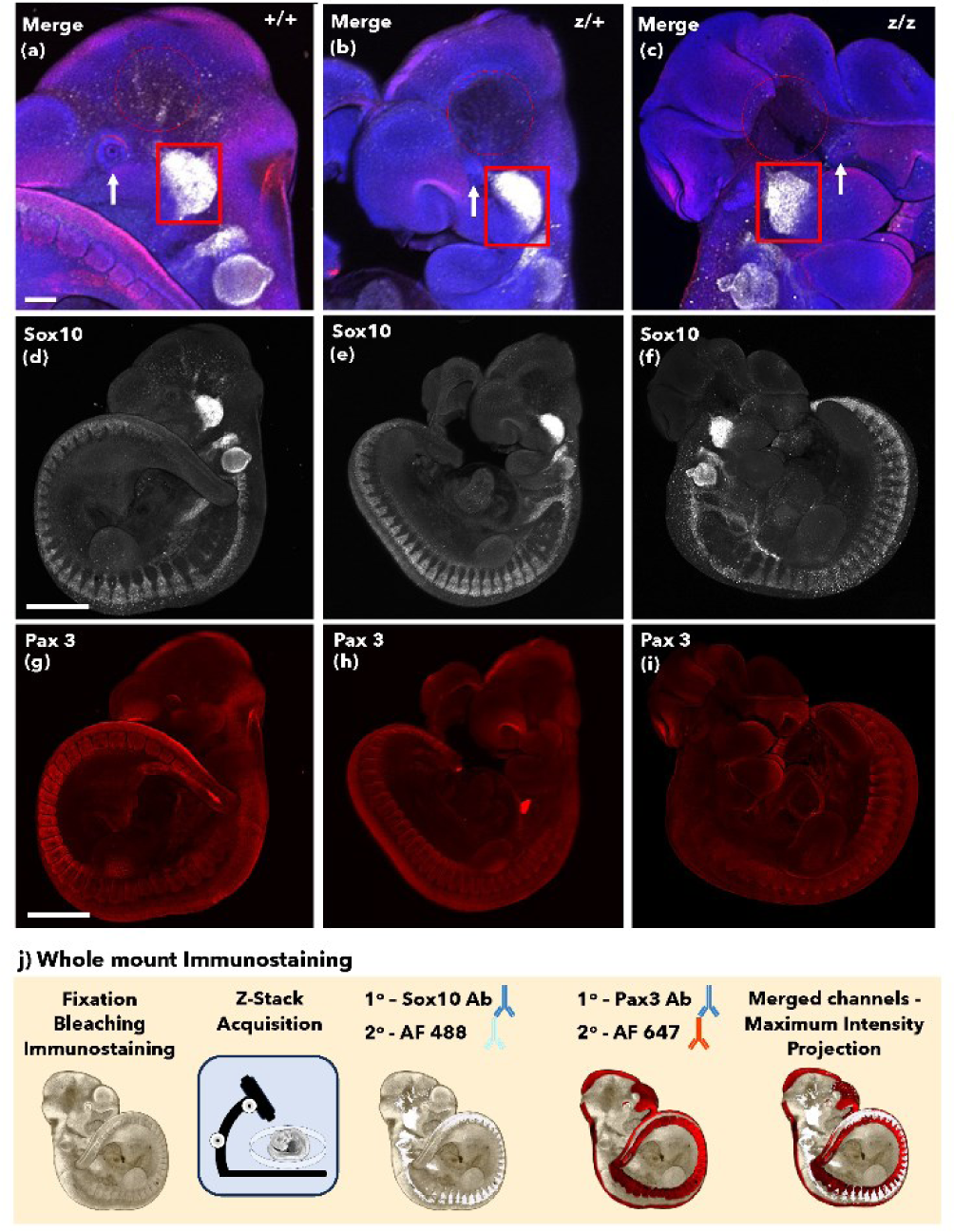
Fuz mutant embryos exhibit abnormal Sox10 positive cranial neural crest migration. Confocal imaging representing the whole mount view of Fuz embryos at E10.5. (a, b, c) Whole mount Sox10 immunostaining; (d, e, f) Whole mount Pax3 immunostaining; (g, h, i) Cranial view of Pax3 and Sox10 merged channels. (j) Illustration of the whole mount immunostaining protocol and image acquisition on a confocal microscope. The scale bar is 0.5 mm.

## DISCUSSION

Primary neurulation begins when the ectoderm gives rise to a flat sheet of cells forming a neural plate. The midline cells along the neural plate will then express Planar Cell Polarity (PCP) genes to coordinate convergent and extension movements, following a cranial-caudal orientation, which makes them form an elongated, but at same time narrower neural groove. Consistent with observations reported in previous studies (Gray *et al*., 2009; Zhang *et al*., 2011; Tabler *et al*., 2013) the Fuz ablated embryos do not present with craniorachischisis, which is the characteristic phenotype observed in mutant mouse embryos for PCP regulatory genes (*Celsr1; Dact1; Dvl1; Dvl2; Dvl3; Ptk7; Scrib; Vangl1; Vangl2*). Instead, *Fuz* ablated embryos present with exencephaly and/or spina bifida, which is associated with mutant mouse models for genes involved in ciliogenesis or transduction of Sonic Hedgehog signaling (*Fuz, Intu, Gli3, Gpr161, Rab23 and Sufu*). Our observations of *Fuz* ablated mouse embryos during hindbrain development at E9.5 and E10.5, indicate that the transient division of the hindbrain, defining eight transverse segments termed rhombomeres (r1 to r8) is abnormal, distinguished by diminished length and width between rhombomere borders (Figure 1 a-i). This observation suggests a hypoplastic development of the hindbrain in the absence of Fuz, where the unique molecular cues expressed by each rhombomere would be disturbed, disrupting the generation of cranial nerves and also the spatial organization of neural crest cell migration.

In most vertebrate models of neurulation, ventral sources of Sonic Hedgehog signaling stimulate the proliferation of the left and right neural progenitors, elevating two apposed neuroepithelial folds from the floor plate. The specification of the roof plate is then mainly driven by BMP signaling inhibition and Wnt signaling activation, until the two layers of apposed neuroepithelia grow enough to reach each other dorsally, forming closure points denominated neuropores. As the notochord provides the initial source of secreted Shh, the neural fold cells positioned ventrally will be exposed to higher concentrations of Shh than the dorsally located cells, establishing a morphogenetic gradient during neural fold growth (Persson *et al*., 2002; Murdoch & Copp, 2010; Zhang *et al*., 2011).

To comprehend how biomechanical forces can be altered on the verge of causing neural tube defects, it is fundamental to develop technologies with enough capability to measure and map mechanical forces guiding morphogenesis. The most traditional techniques providing high-resolution maps, such as atomic force microscopy, magnetic-bead twisting, optical tweezers, and micropipette aspiration (Bao and Suresh, 2003), require contact and therefore are limited for live measurements. Non-contact methods, including acoustic microscopy (Quate *et al.,* 1979) or elastography (Muthupillai *et al.,* 1995), regardless of great advances in optical coherence elastography technology (Wang and Larin, 2015; Larin, and Sampson, 2017), are limited for embryonic studies due to their lack of spatial resolution and mechanical accuracy.

The images represented in Figures 2 and 3 exhibit the use of a novel, multimodal imaging technique combining high resolution optical coherence tomography (OCT) and Brillouin microscopy (Ambekar *et al*., 2022) to evaluate tissue structure and stiffness during hindbrain neural fold development in *Fuz* ablated embryos. In advance of real-time Brillouin imaging, OCT was used to detect three-dimensional details (∼10 µm spatial resolution) along the rhombomere 5 (r5) and the otic pits (auditory pit), defining a transverse axis to measure tissue stiffness based on the frequency shift of Brillouin light scattering (GHz). Our data shows that neuroepithelial stiffness is decreased in the absence of Fuz at E9.5 (Figure 2), while the most dorsal areas of the neural folds partially recover stiffness at E10.5, indicating persistent loss of ventral neural fold stiffness throughout neurulation.

Previous studies in mouse embryos have shown that ablation of genes with a function upstream of Shh signaling pathway reduces the strength of Shh signaling, being associated with the midline defect holoprosencephaly (Murdoch and Copp, 2010), including *Shh, Smo, Stil, Hhat, Cdon* and *Gas1* (MP:0005157), while ablation of genes acting downstream of Shh are associated with exencephaly and spina bifida (MP:0000914 and MP:0003054), and respective loss of Shh inhibitory influences that can lead to an increase in Shh signaling, including *Gli3, Gpr161, Rab23* and *Sufu*, which are mouse phenotypes also observed after ablation of genes required for the last steps during ciliogenesis (*Dync2i2, Fuz, Ift122, Ift140, Ift57, Intu* and *Kif7*).

Our stiffness measurements at E9.5 and E10.5 are pointing towards a direction where impaired cilia development, in the absence of Fuz, disturbs transduction of Sonic Hedgehog signaling and as a consequence, the neuroepithelial floor plate does not develop properly, confirming our initial observation of a hypoplastic hindbrain phenotype in this mouse line (Figures 1, 2 and 3).

When studying embryonic mouse mutants is important to perceive that mid-gestation lethality can mask NTDs, and careful observation is needed to ensure that a certain phenotype is responsible for disturbing normal neurulation (Copp *et al*., 2003). Neuroepithelial cells rely on primary cilium for transduction of Shh signaling, which is a requirement to maintain neuroepithelial cells as cycling neural progenitors. During neuronal differentiation, the cell bodies of neuroepithelial cells are translocated toward their basal poles until they delaminate from the neuroepithelium, in a process involving loss of apical polarity and primary cilium disassembly (Das and Storey, 2014). Our observations using Pax6 and Tubb3 staining to map neuronal differentiation indicate that normal development of the cranial and paravertebral ganglia is impaired in Fuz mutant embryos, and the pattern of differentiated neurons along the body axis is significantly decreased, indicating that the function of Fuz in neuroepithelial cells is required for neurogenesis (Figure 4).

Immunostaining for Sox10 positive neural crest cells also indicate impaired development of cranial ganglia and paravertebral ganglia, including more posterior areas along the rostral caudal axis (Figure 5). Cranial neural crest cells contribute to the cranial ganglia, primarily delaminating from the even numbered rhombomeres 2 and 4 towards the first and second pharyngeal arches, respectively, while the population of neural crest cells derived from the rhombomeres 6 until 8 will migrate posteriorly, towards the third to sixth pharyngeal arches (Serbedzija *et al*., 1992; Sechrist *et al*., 1993; Inman *et al*., 2018). Considering the hypoplastic hindbrain development in *Fuz* ablated embryos, presenting abnormal rhombomere segmentation, we have also performed Pax3 immunostaining. The normal Pax3 pattern is disrupted at the midbrain-hindbrain boundary in the *Fuz* ablated embryos, indicating abnormalities during isthmic organizer formation (Figure 5).

As the neural plate elongates during neurulation, the diencephalon (forebrain), mesencephalon (midbrain), and rhombencephalon (hindbrain) are defined anteriorly at the neuraxis. The boundary established between the mesencephalon and rhombencephalon forms the isthmic organizer, a transient signaling center, which works to regulate the development of anatomical structures derived from the midbrain-hindbrain boundaries. Malformations of the isthmic organizer lead to anterior NTDs, including exencephaly, a severe type of NTD in which the cranial neural folds are overturned, also presenting unfused skull bones, which results in brain exposure and early lethality (Copp, 2005). Taken together, this work shows for the first time that hypoplastic hindbrain development, presenting abnormal rhombomere morphology and persistent loss of ventral neuroepithelial stiffness precedes exencephaly in Fuz ablated embryos.

## MATERIALS AND METHODS

### Animal Husbandry and Mice Timed Mating

All mice were housed and handled in accordance with Institutional Animal Care and Use Committees approved protocols at the Baylor College of Medicine and the University of Houston. To obtain embryos at specific developmental stages, a controlled breeding strategy was used to select the breeding pairs, to introduce the male and female in the same cage, to observe the presence of a copulatory plug, to keep track of the pregnant female and collect the embryos at the correct embryonic day. Embryos were dissected on gestational days E9.5 and E10.5 and kept in 100% rat serum at Baylor College of Medicine. The fresh embryos were then transferred to the University of Houston within 30 minutes. The yolk sac was removed for genotyping, and the embryos were mounted to expose the hindbrain neuropore for Brillouin-OCT measurements.

### Multimodal Brillouin-OCT system

The multimodal Brillouin-OCT system has been described in a previous publication (Ambekar *et al*., 2021). Briefly, the home-built system consists of swept source OCT sub-system, dual VIPA-based Brillouin microscopy sub-system and a combined scanning arm. The Brillouin microscopy system consisted of 660 nm single mode laser source (Torus, Laser Quantum., Inc, CA). The incident power on the sample was 35 mW. The collected back scattered light from the sample was transferred to the dual VIPA spectrometer (Berghaus et al, 2015) and an electron multiplying charge coupled device camera was used to detect the Brillouin frequency shift of the sample. The camera acquisition time was 0.2 s per spectrum acquisition. The system was calibrated with standard materials such as water, acetone, and methanol before every measurement. The sample was imaged with a microscope objective with 0.25 NA, resulting in an axial resolution of ∼36 μm and lateral resolution of ∼3.8 μm. The swept source OCT system had a central wavelength of ∼1310 nm, scan rate of ∼50 kHz, scan range of ∼105 nm, and ∼8 mW incident power on the sample. The lateral and axial resolutions were ∼17.5 μm and ∼10 μm in air, respectively. Light from both systems was combined using a dichroic mirror, and galvanometer-mounted mirrors scanned the beam across the sample. For Brillouin imaging, the sample was stepped by a motorized vertical stage. A custom instrumentation software was developed that utilizes the OCT structural image to guide Brillouin imaging.

### Immunohistochemistry

Pax6 (42-6600, ThermoFisher) and Tubb3 (2H3-Tuj1, DSHB); Sox10 and Pax3 (DSHB) whole-mount immunostaining was performed on E10.5 embryos, after fixation for 30 minutes in 4% PFA. To remove the fixative agent, the embryos were washed three times in PBS for five minutes, and a consecutive dehydration step to 100% methanol. Endogenous peroxidase activity was blocked by incubation in Dent’s bleach (MeOH:DMSO:30%H_2_O_2_; 4:1:1) for 1 hour at room temperature. Embryos were rehydrated in TBS containing 0.1% Tween-20 (TBST) for 30 minutes and incubated in a 1:500 dilution of anti-Pax6 and anti-Tubb3 overnight at room temperature. Embryos were washed five times in PBST for five minutes, and then incubated overnight at room temperature, with a 1:1000 dilution of fluorescent secondary antibodies in TBST. On the next day, the embryos were washed five times in TBST, for five minutes. Before general cell visualization, the embryos were incubated overnight in DAPI (4′,6-diamidine-2-phenylidole-dihydrochloride; 2 µg/ml) to label all the nuclei. Before visualization on the microscope, the embryos were cleared in 25% glycerol and imaged using a Nikon CSU-W1 Yokogawa spinning disc confocal. The obtained Z-stacks were projected at maximum intensity using the Nikon NIS-Elements software and exported as TIFF files (Inman *et al*., 2018).

### Statistics

For statistical analysis, significance levels were assessed using One-Way ANOVA and Bonferroni’s comparisons test. * and **** indicate P < 0.05 and P <0.0001, respectively. Differences were considered statistically significant at * and **** indicate P < 0.05 and P <0.0001. All calculations were performed using the GraphPad Prism Software.

## ACKNOWLEDGEMENTS

The authors would like to thank the Baylor College of Medicine core facilities for their important contributions in the beginning of this work, particularly the Baylor College of Medicine Integrated Microscopy Core Facility.

## COMPETING INTERESTS

Drs. Finnell and Wlodarczyk participated in TeratOmic Consulting LLC, a now defunct consulting company. Additionally, Dr. Finnell serves on the editorial board for the journal Reproductive and Developmental Medicine and receives travel funds to attend editorial board meetings. Drs. Singh and Larin have a financial interest in ElastEye LLC., which is not related to this work. All other authors have no conflict of interest to declare.

## FUNDING

This project was supported by grants from National Institutes of Health (R01 HD095520 to K.L., G.S. and R.H.F.) and National Science Foundation (DBI-1942003 to G.S.).

## AUTHOR CONTRIBUTIONS STATEMENT (CRediT taxonomy)

**C.D.C.**, Conceptualization, Formal Analysis, Investigation, Methodology, Resources, Visualization, Writing – original draft, Writing – review & editing

**Y.S.A.,** Conceptualization, Formal Analysis, Investigation, Methodology, Visualization, Writing – review & editing

**M.S.,** Formal Analysis, Methodology, Methodology, Visualization

**L.Y.L.,** Resources

**B.W.,** Resources

**S.R.A.,** Supervision, Formal Analysis, Methodology, Visualization, Writing – review & editing

**G.S.,** Supervision, Writing – review & editing, Project administration, Funding acquisition

**K.V.L.,** Supervision, Writing – review & editing, Project administration, Funding acquisition

**R.H.F.,** Supervision, Writing – review & editing, Project administration, Funding acquisition

